# Awareness of the future: Dolphins know when they need to remember for the future

**DOI:** 10.1101/2024.11.28.625857

**Authors:** Sara Torres Ortiz, Simeon Q. Smeele, Mathias Osvath, Andrea Martín Guerrero, Ariana Hernandez Sanchez, Cristina Ubero Ramirez, Javier Almunia, Auguste M. P. von Bayern

## Abstract

In humans, awareness of an upcoming memory test enhances intentional encoding and improves memory recall. Here, we investigated whether dolphins exhibit similar future-oriented encoding of information known to be needed in the future. Dolphins were trained to remember specific, randomly assigned actions for later re-enactment, with either immediate or delayed recall. When an unexpected delay was introduced in trials anticipating immediate recall, memory was retained for only 13 seconds, suggesting working memory encoding. However, when instructed to expect delayed recall, dolphins accurately reproduced actions after delays even after 16 hours. These results suggest that dolphins, anticipating future need, intentionally encode actions to be performed in the future into long-term memory, implying prospective encoding and prospective memory capacities. Their memory also displayed key features of episodic memory: encoding occurred in a single episode, and memory was declarative, as the action itself declared its content. Moreover, dolphins more effectively recalled self-performed actions compared to gestural codifications of the same actions, mirroring the human-typical “enactment effect” and supporting episodic-like memory over semantic memory. Our findings indicate that dolphins show awareness of future memory demands and seem to use a future-oriented, episodic-like memory system, capable of storing prospectively encoded, intended actions in long-term memory.

## Introduction

Memory is a fundamental cognitive process that allows organisms to store and recall information about past experiences. Among humans, one of the most advanced forms of memory is episodic memory, which involves recalling specific events within their temporal and spatial context ^1^. A crucial subset of episodic memory is prospective memory—the ability to remember to perform an intended action at a future time ^2^. The ability to plan and act with the future in mind is essential for many aspects of daily life, from remembering to take medication, booking train tickets, preserving food supplies for winter to anticipating future needs in general.

Human memory studies have shown that people are often more likely to remember information when they are aware that it will be needed in the future ^3–5^. For example, adults demonstrated better memory performance when they knew beforehand that they would be tested on an episodic memory task—specifically, recalling which items were hidden, in which location, and during which session—compared to when they were asked to recall the same information incidentally, without prior knowledge of its relevance to a later memory test^3,4,6^. In such studies, participants who know they will need to recall specific information later are more likely to intentionally encode that information into working memory ^5,7^ but also long-term memory ^4^. In contrast, when information was perceived as only temporarily relevant, it is often forgotten once it is no longer needed ^8,9^.

While research on humans has explored how the awareness of future demands influences memory encoding^10,11^, this has not been studied in non-human animals. The concept of “intentional encoding” as it is understood in humans—where participants are explicitly aware that they need to remember information for the future and can articulate that awareness—is difficult to translate directly to animal research due to communication barriers and the challenge of assessing animals’ awareness of their own memory processes ^12–14^. Most of the research on animal memory has focused on retrospective memory—recalling past events or learned information (e.g. ^12,15,16^). The few investigations into prospective memory, have focused on appropriately-timed recall (of typically one specific target action) rather than the intentional encoding of information for future use. This gap in research leaves unanswered questions about the extent to which non-human animals share the human ability to plan and remember future intentions.

In this study, we aimed to address this gap by exploring whether dolphins, which stand out among vertebrates for their large relative brain size and cognitive performance ^17^, demonstrate this aspect of prospective memory. Specifically, we investigated whether dolphins can intentionally encode actions to be recalled and performed after varying delays, thereby showing an awareness of future needs. This approach not only contributes to our understanding of dolphin cognition but also offers insights into the evolutionary origins of future prospection. Specifically, to investigate whether dolphins exhibit intentional encoding for future use, we designed a series of experiments that took advantage of their trainability, employing two commands specifically trained for this study.

In the first experiment, we focused on memory for own actions^18^ when the dolphins anticipated immediate recall. Dolphins participated in interactive sessions with a trainer, where they were asked to perform arbitrary trained actions on command in random order. In 54% of trials, they were required to “repeat” the last action they had performed (using a “repeat command”), whereas in 46% of trials they were instructed to perform another trained action. This setup allowed us to test whether dolphins could retain the memory of an action over a short period, anticipating the possibility of needing to repeat it. Hence, after a short interval of 5 seconds, they either had to retrieve their working memory or update it. Once the dolphin’s baseline performance was established, we introduced an unexpected delay.

The second experiment was designed to explore whether dolphins could encode actions for extended retention intervals based on an anticipated future need (i.e., anticipated recall after a delay). Here, dolphins were trained to respond to a “remember” command, which instructed them to encode a trained action for future reenactment after delays of up to 15 minutes before testing them in increasing delays. We hypothesized that if dolphins could intentionally encode actions into long-term memory when they understood they would need to recall them later, they would demonstrate successful memory recall even after extended delays. In contrast, we expected memory retention to decline rapidly in the condition in which subjects anticipated immediate recall.

Finally, we examined whether enacting the action to be remembered during encoding improved memory performance compared to simply receiving the corresponding gestural command. We hypothesized that if enacting the action enhanced memory, it would suggest that dolphins form an intention to reenact the previously self-performed behavior, rather than relying on semantic memory.

## Results

### Experiment 1: Anticipating immediate recall, dolphins store their last self-performed actions in working memory only

The first experiment examined the memory capabilities of dolphins (LP dolphins, 0’-delay condition) when they expected either having to repeat a previous action a few seconds later or having to perform and remember another action hence updating their memory. Three dolphins were tested using the “repeat” paradigm established by Mercado ^18^, by which he demonstrated that dolphins can recall and reenact their recent own actions from working memory. Initially, the dolphins went through an acquisition phase to learn the abstract rule of repeating their last action on command (“repeat” command, see Methods). In the subsequent experiment, they either had to perform one out of four actions on command in random order (action trials, 46%), or, within three seconds, were asked to repeat the previous action by the “repeat command” (repeat trials, 46%) (see Fig 1A).

**Fig 1.**
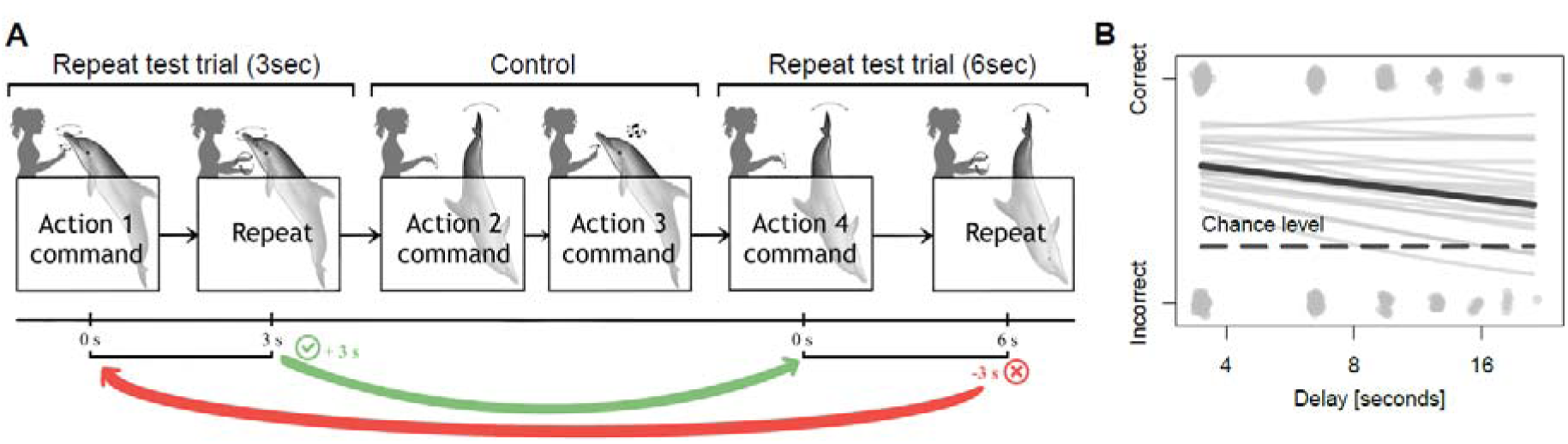
Memory performance when anticipating immediate recall of own actions (Experiment 1). (**A**) Illustration showing a series of gestural commands given by the trainers in three exemplary consecutive trials (two repeat trials and one interspersed action trial are shown). If the dolphin performs correctly in a repeat trial, the delay between the “action command” and the “repeat command” increased 3 seconds in the subsequent test trial (see green arrow). If the response in a test trial was incorrect, the delay decreased 3 seconds again (see red arrow). (**B**) Memory performance when unexpected 3-second delays were introduced incrementally. The delay between the performed action and the “repeat command” is shown on the x-axis (in seconds). The dolphins’ response is indicated on the y-axis. The thick black line denotes model average predictions. Grey dots represent outcomes of trials (correct or incorrect) across all three individuals. Each grey line summarises the samples of the posterior. The dashed black line represents chance-level (1/4).

All three dolphins successfully repeated their previous actions when there was no delay (average performance across 208 trials/animal: 64%; 89% PI: 49%, 76%). However, when delays were introduced in 3-second increments, the dolphins could only recall for 13 seconds (Fig 1B; S1 Video). Beyond this point, their performance dropped to chance level.

### Experiment 2: Anticipating recall after a delay, dolphins explicitly encode self-performed actions in long-term memory

Five dolphins (DA dolphins, 15’-delay condition) underwent an acquisition phase to learn the abstract rule to remember specific self-performed actions on command (“remember” command, see Fig 2A) following progressively longer delays, starting from one minute and extending up to 15 minutes. In contrast, two dolphins (LP dolphins) in a 3’-delay condition never experienc d delays exceeding three minutes during the acquisition phase (see below). This preparation allow d the dolphins to anticipate “recall requests” after prolonged or brief intervals (refer to Fig 2A; 2 Video and SI for details). Our aim was to investigate whether the instruction to remember (“remember” command) indeed functioned as a cue for a future “need” of the memory, leading to encoding into long-term memory and whether the expected recall delay influenced the memory.

**Fig 2.**
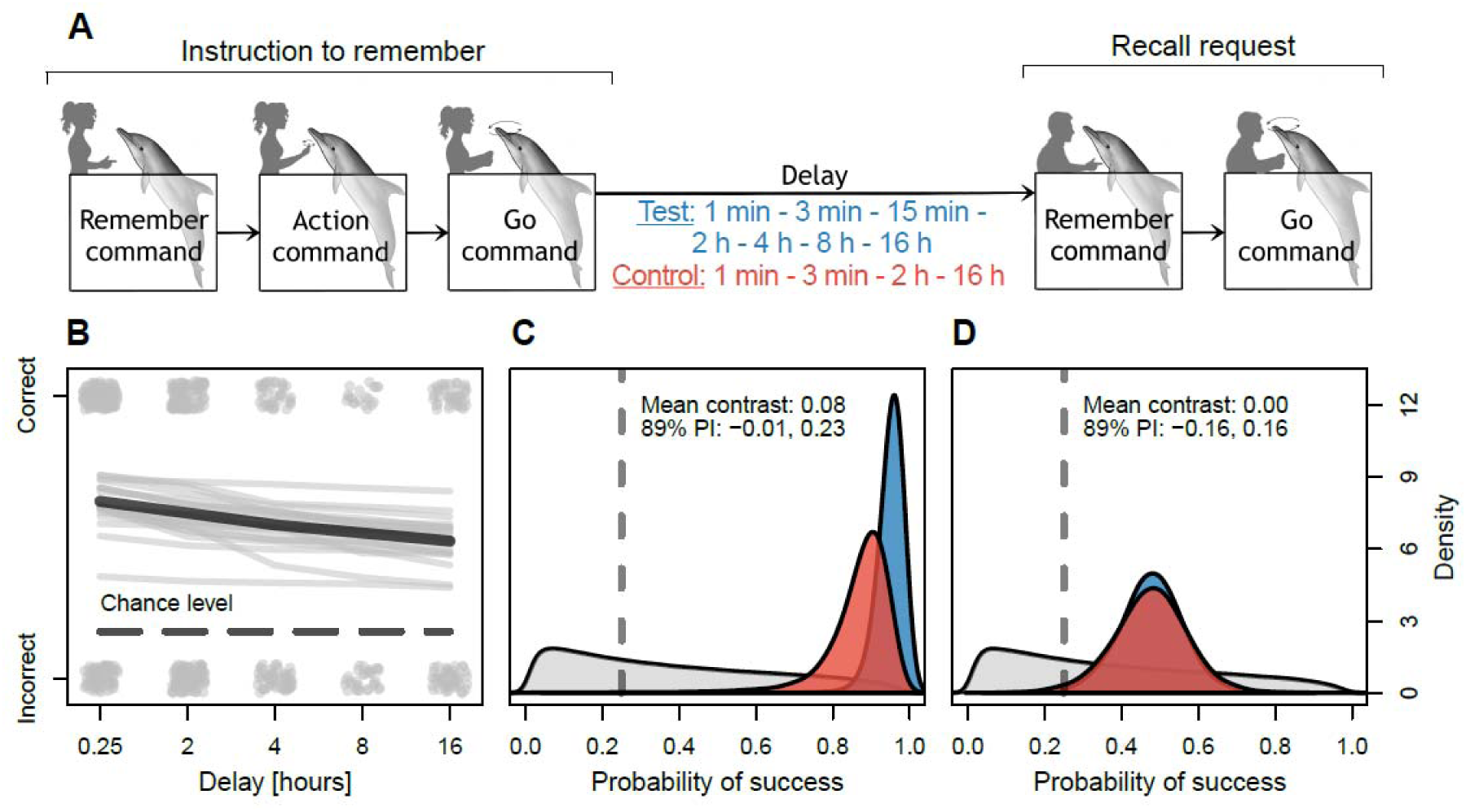
Memory performance when anticipating recall of own actions after long delays (Experiment 2). (**A**) Series of commands given by the trainers in each trial. The first three commands given by one trainer instruct the dolphin to perform and remember a specific action (“remember command”). After the delay, a different trainer, naive of the action to be performed, requested the retrieval of the remembered action (“recall command”) by two consecutive commands (“remember” and “go”; see delay implementation in red and blue letters and Methods). (**B**) Memory performance of the 15’-delay group. The recall delay for retrieving the remembered action is shown on the x-axis (in hours). The dolphins’ response is indicated on the y-axis. Grey dots represent outcomes of trials (correct or incorrect) across all five individuals. The thick black line denotes model average predictions across phases. Thin grey lines show 20 samples of the posterior. The dashed black line represents chance-level (1/6). (**C & D**) Gray density plots show the prior centred around chance-level (grey dashed line). Blue coloured density plots show the posterior distributions for the average performance of the 15’-delay group and red coloured density plots represent the 3’-delay group for (**C**) the baseline trials (3 minutes delay) and for (**D**) the 2 hour delay test phase.

When subsequently tested, the 15’-delay condition dolphins explicitly encoded and successfully recalled specific actions they had performed upon trainer instruction up to 16 hours later, the maximum duration tested (see methods and SI for specifics; see Fig 2B; average performance at 16 hours: 58%; 89% PI: 37%, 75%). To investigate the effect of distractions, half of the tests were embedded in the ongoing daily activities in the dolphinarium. Hence, the subjects took part in various training sessions and interacted with different trainers throughout their delay times, performing up to 251 actions on trainer command during the 8-hour delay (see SI). The dolphins’ memory performance was neither meaningfully affected by increasing delay duration (β_phase_: 0.57; 89% PI: 0.14, 1.02) nor by distracting activities occurring during the delay (β_distractions_: 0.16; 89% PI: −0.23, 0.55).

The two dolphins in the 3’-delay group had been equally trained to acquire the “remember” and “recall” commands, however, they had never encountered a recall delay exceeding three minutes before being subjected to a 2-hour delay test. Thus, they were not anticipating a recall test occurring much later. Nonetheless, they performed comparably to the 15’-delay group, which had prior long-delay training, both during the baseline (recall command after a 60-second delay) and the two-hour delay test (Average log-odds difference between the 15’-delay group and 3’-delay group for the 60-second delay: −1.10; 89% PI = −2.70, 0.14; for the 2-hour delay: −0.01; 89% PI = −0.66, 0.64; Figs 2A, C and D, and SI). The two 3’-delay group dolphins also spontaneously recalled successfully following an even longer unexpected delay trial after 16-17 hours.

### Experiment 3: Enactment effect: dolphins remember a to-be-performed action better after having previously self-performed it

To assess whether performing the action, instead of just being exposed to the command for the action, would impact on the dolphin’s memory, the two 3’-delay condition dolphins underwent an additional experiment. Instead of the “remember” command as shown in Fig 2A, the subjects were given the “remember” and “action” commands (Fig 3A). However, the command to perform the indicated action (“go” command) was delayed (10 seconds and 60 seconds). Therefore, the subject was instructed which action to remember for future enactment but could not perform it until after the delay (Fig 3A; S3 Video).

**Fig 3.**
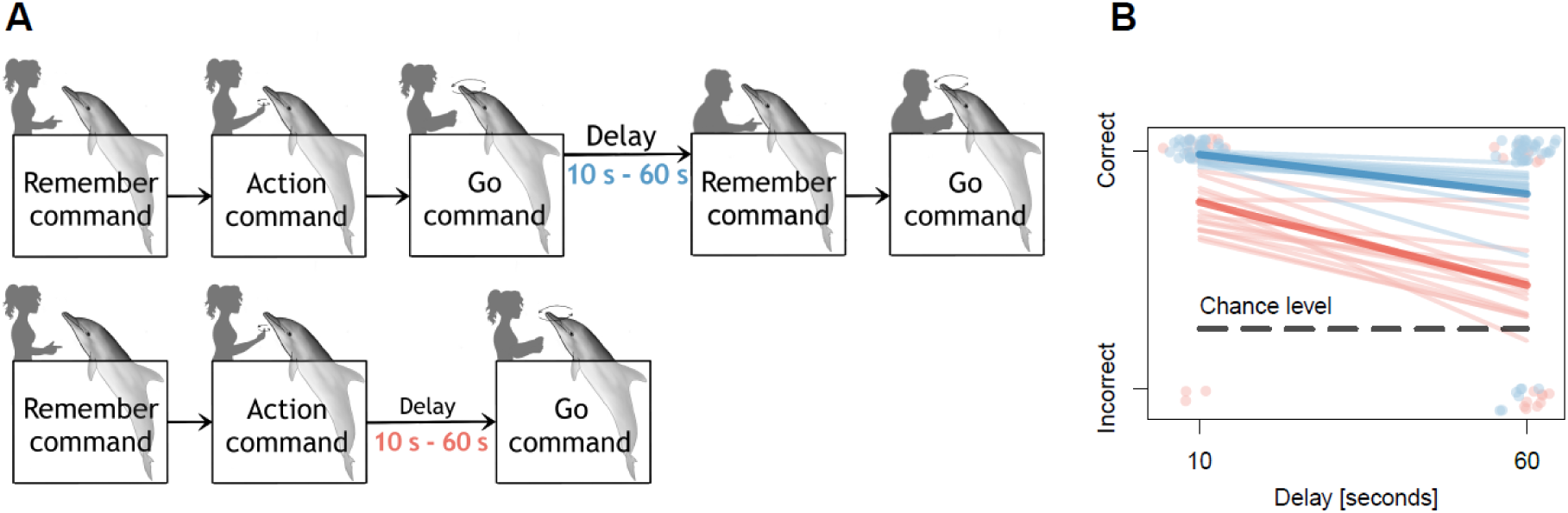
Comparison of the memory performance of previously enacted versus gesturally encoded actions (Experiment 3). (**A**) Illustration showing the series of gestural commands given by the trainers in each trial. The first two commands instruct the dolphin to remember a specific action. After the delay, the trainer requested memory recall by giving the “go command”. (**B**) Enactment effect: memory performance decreases when remembering gesturally encoded, previously not self-performed actions compared to previously self-performed actions. The graph shows a comparison of the memory performance when recalling a previously self-performed action (=baseline in blue) and a not self-performed but gesturally encoded action (red) after a 10 second and a 60 second delay. The dolphins’ response is indicated on the y-axis. Coloured dots represent outcomes of trials (correct or incorrect) across both subjects and the thick lines denote model average predictions. Each thin coloured line illustrates a sample from the posterior. The dashed black line represents chance-level (1/4).

With a 10-second delay between “action” command and “go” command, the dolphins correctly performed the action in 6 out of 8 trials. However, with a 60-second delay, their performance dropped significantly to 3 correct trials out of 8 (average contrast (log-odds): −1.55; 89% PI: −2.9, −0.29). Compared directly to the 60-second delay, the dolphins performed significantly better in baseline trials (Fig 3B), in which they had to remember and reenact a previously self-perform d action after 60 seconds (average contrast (log-odds): 1.76; 89% PI: 0.73, 2.82) than in the 0 seconds delay trials without prior enactment

## Discussion

Our study shows that dolphins recognize future need to remember. When anticipating a future memory test, they intentionally encode information for future use, comparable to humans ^3–5^, and recall even after long retention times. Depending on whether they expected the memory test in the more distant or immediate future, the dolphins explicitly encoded it in long-term or working memory respectively, suggesting memory control ^19^ and corroborating their awareness of the future demand. When anticipating immediate recall (or only very short delays never exceeded five seconds) in the “repeat” context (0’-delay condition), the dolphins stored their own actions in working memory and therefore failed to remember after unexpected delays exceeding 13 seconds. This may indicate a working memory limit ^20^ very similar to those of other vertebrate species tested with the “repeat” paradigm ^21–23^ but could also represent a case of directed forgetting ^24^. According to their prior experience in the repeat context, in which they were either requested to recall within five seconds or to update their memory, the stored information became irrelevant after short delays. (Directed) forgetting of information not considered relevant any longer stored either in working or long-term memory is well-studied in humans ^8,9^ and also described in animals (see ^25^ for review). When however, the dolphins had been instructed beforehand (using the “remember” command) that the action had to be remembered for re-enactment in the more distant future, they intentionally encoded it in long-term memory, likely as an intended action with motor cortex involvement (see below). They were able to reenact the remembered action successfully after delays up to 16 hours (the maximum delay tested), which is a retention time of a magnitude previously only shown in dogs for remembering a specific action that was demonstrated by a human experimenter ^26^.

The dolphins’ intentional encoding of actions into long-term memory for future enacting, is suggestive of prospective memory ^27^ and indicates so-called “prospective encoding”, i.e., the encoding of a future response for later implementation ^28^. Prospective coding is considered a component of prospective memory and suggests the ability to represent future events ^15^. Evidence for prospective coding across various amniotic endotherm species, from pigeons ^29–31^ to rats ^32^ and primates ^33,34^ indicates that it may be a feature of the mammalian neocortex and of the pallial homologue in birds, making its presence in dolphins expected. Recognition of the future need to recall and explicit encoding, as our study reports for dolphins, are prerequisites of prospective memory ^27^. Consistent with prospective memory, we can show that their memory was not affected by distractions, which would have interrupted any rehearsing, nor by increasing delay duration. The dolphins therefore must have stored the information in long-term memory rather than continuously rehearsing it ^35,36^. Concerning the recall, a key aspect of prospective memory is that the retrieval of a stored intention should occur spontaneously in the appropriate moment, without external prompting ^35^. One may argue that the subjects in this study were prompted to recall by the trainer and did not themselves remember to act at a particular moment. However, several studies reporting event-based prospective memory in humans in fact used verbal recall cues very similar to the neutral recall requests made by the dolphin trainers in our study ^37–39^. In any case, the dolphins reliably performed the action they were asked to remember, which constitutes the “retrospective component” of a prospective memory task ^35^.

The finding that the dolphins in the 3’-delay condition recalled the encoded action when spontaneously tested with never previously encountered long delays (2 hours and even 16-17 hours), demonstrates that when a future need exceeding working memory capacity (presumably about 15 seconds) is anticipated the action, is intentionally encoded in long-term declarative memory. Human declarative memory refers to memories that can be “declared” ^40^ and comprises both semantic memory (general knowledge and facts) and episodic memory (remembering personally experienced events) ^41,42^. The dolphins in our study could declare their memory content by enacting it, and hence exhibited declarative long-term memories of their own actions. Given that they were encoded in a single episode (single-trial learning), our findings cover several key features of human episodic memory ^16,43,44^. Episodic memory involves re-experiencing specific self-experienced situations in the past through subjective mental time travel ^16^, which typically constitutes recalling from a first-person perspective what one (or others) did in a particular past situation. Remembering self-performed actions is special in this regard because the remembered action both constitutes the self-experienced past episode and declares the memory content.

Another key finding of our study was that the dolphins showed an “enactment effect” similar to that reported in humans ^45,46^. In humans, memory research has revealed that memory performance is better when actions are enacted during encoding compared to standard verbal encoding (i.e., reading or hearing verbal instructions to be remembered ^47,48^). Similarly, the dolphins could remember better when they re-enacted an action which they had previously self-performed than when they had to perform an action that previously just had been indicated abstractly to them, by gestural command. This finding implies that there is something fundamentally different about encoding own actions ^49^. In humans imaging studies demonstrate that enacting during encoding leads to motor cortex reactivation when recalling ^50,51^, and lesion studies confirm the involvement of movement representations in encoding episodic memories of action ^52^. It is therefore plausible that also the motor cortex of dolphins is activated both during encoding and during recall, which supports that they are re-experiencing their prior self-performed action and that their memory of own actions episodic rather than semantic. If the dolphin’s memory was purely semantic, no such self-performance dependent enactment effect would have been expected.

Interestingly, recent discoveries show that the enactment effect in humans extends to intended actions ^53,54^, making it additionally relevant for the study of prospective memory ^55^. In humans, recall of intended actions is typically excellent ^56^ while prospective memory failures occur due to mistiming rather than forgetting intentions ^2^. This has been linked to the fact that delayed intentions involve actions to be enacted later ^56,57^ implying preparatory motor operations during encoding, which is supported by experimental and brain imaging studies ^56^. This highlights the differences between prospective memory (to-be-performed tasks) and retrospective memory (to-be-remembered tasks) not only in retrieval but also in encoding processes and storage characteristics ^56^. The discovery of the enactment effect in dolphins, and in the context of prospective memory, prompts further investigation to determine if similar processes exist in distantly related mammals.

A possible evolutionary explanation for the dolphins’ prospective and episodic memory abilities relate to their complex socio-ecological environment and behaviour. Dolphins live in highly complex fission-fusion societies ^58^ characterized by flexible group sizes that can reach up to 100 individuals ^59,60^, and form diverse social alliances and long-term affiliative bonds ^61,62^. Moreover, they display collaborative hunting ^63^, the most complex form of hunting where different individuals acquire and maintain different roles throughout the hunt to achieve a joint goal ^64,65^, which arguably is future-oriented and may require basic planning abilities. Forming prospective memories and remembering one’s past interactions with specific conspecifics may help these intricately social animals to optimize their behaviour in highly dynamic fission-fusion societies, avoiding social conflict situations, finding suitable hunting alliances or determine who to associate or ally with and whom to avoid.^61,66^.

Dolphins, as aquatic mammals, boast remarkable anatomical, sensory, and behavioural adaptations^64,67–69^. They starkly differ in appearance and lifestyle from terrestrial mammals, which have evolved over hundreds of millions of years on land. Despite this distinction, dolphin brains exhibit typical mammalian traits along with superior cognitive abilities ^62,70^ and high neuron-densities ^71^. Our study contributes to the mounting evidence suggesting that episodic memories likely represent a feature of the mammalian brain, relying on shared neuroanatomical features absent in reptiles. For instance, two endotherm lineages convergently evolved about 20 times more neurons than similarly sized ectotherms ^72^, many of which are dedicated to sensory and motor processing in the neocortex. These neurons project to the hippocampus, which has evolved a distinct role compared to non-avian reptiles, facilitating the formation of episodic memories ^73^.

## Conclusion

Our study reveals that dolphins have the capacity to recognize when information will be needed in the immediate or more distant future, intentionally encoding actions into long-term memory when necessary. This ability suggests a form of prospective memory in dolphins, and we further show that they possess declarative long-term memories of their own actions, formed in single episodes—key components of human episodic memory ^16,44,74^.

Additionally, our findings include evidence of an enactment effect, previously documented only in humans, supporting the existence of an episodic memory system for self-initiated actions in dolphins and potentially in other mammals. These results point to the possibility of a shared memory bias among mammals, which aligns with the action-simulation capacities of the mammalian neocortex ^73^.

Future research could expand on these findings by exploring episodic memory and prospective memory across diverse species, using behavioural training and gestural communication as investigative tools, to better understand the evolutionary pathways and cognitive drivers behind this memory bias.

## Materials and methods

### Data and code availability

All original code has been deposited at Zenodo and is publicly available as of the date of publication. The data can be accessed now at the following link: https://github.com/simeonqs/Need_to_remember_Dolphins_recall_their_own_actions_after_long_delays.

### Experimental model and study participant detail

Five female and three male bottlenose dolphins participated in this study. The five 15’-delay condition dolphins were kept in Dolphin Adventure, Puerto Vallarta Mexico (DA dolphins). The subsequently tested three 0’- and 3’-delay condition dolphins were kept in Loro Parque: Animal Embassy, Tenerife, Spain (LP dolphins). The four DA dolphin females in the 15’-delay group, Karina, Eva, Nouba and Lluvia were 30, 10, 19 and 14 years old respectively. Eva and Lluvia were sisters, the other females were not related to each other. These four females were housed in a pool system together with 12 more dolphins in a 4 m deep pool system of 2465 m^2^ (S1 Fig). Eva had been pregnant and Nouba had a calf lactating during the entire duration of the experiment. Karina was wild-born, and Eva, Nouba and Lluvia were captive-born in their housing facility, and they were 30, 10, 19 and 14 years old, respectively. The DA dolphin male in the 15’-delay group, Nemo, was 15 years old and captive-born in Dolphin adventure. He was housed in a different pool system (1373 m^2^; 4 m deep; See S2 Fig) than the females, together with six other dolphins (S2 Fig).

The three 0’-delay group dolphins from Loro Parque, Achille, Ulisse and Clara were captive-born. Clara was born in Loro Parque and was 23 years old during the experiment while Achille and Ulisse were born in an Italian dolphin facility and were group-housed together with six other dolphins in a system of pools (S3 Fig) with interconnected pools between which the animals rotated every day (max. depth 6 m and 7 million litre). The pools were outdoors and exposed to natural weather and light conditions. Ulisse was 25 and Achille was 20 years old when the experiment took place. During training and the test, Achille and Ulisse stayed together while Clara was in a separate pool as she had a one-year-old calf with her.

All dolphins had 4-5 training (or scientific testing) sessions throughout their day, during which they were kept active and received their full diet (independent of how well they performed). They were fed with a variety of fish (e.g., a mix of herring, sprat, capelin, and blue whiting). The training sessions were important for the animal’s wellbeing (enrichment) as well as establishing medical procedures. All dolphins participated voluntarily both in the training sessions and the experimental sessions. Food deprivation was never used to increase the dolphins’ motivation (83, 84). All dolphins had been trained for multiple years and thus responded to various standard commands employed by dolphin trainers, important for behaviour training but also for medical procedures (e.g., to come, to stay at a “station”, to turn upside down, to cross the pool, to open their mouth etc.). They also exhibited a large repertoire of different trained actions they performed reliably on specific trainer commands (action commands). For this experiment a subset of four trained actions was chosen.

All subjects were trained or tested individually in their home pool while the remaining group members were also engaged in training sessions running in parallel with different trainers in different locations of the pool. The reason for testing the dolphins in a group set up was that a separation from their group would have produced unnecessary stress for these highly social animals and would have resulted in reduced attention, which could have impacted on their memory. During training, all available fish species were used as rewards. Yet, during our testing sessions, only herring was used so as not to influence the dolphins’ response with varying reinforcement.

### Method details

The dolphin memory study consisted of three experiments. Experiment 1: memory performance when anticipating immediate recall, Experiment 2: memory performance when anticipating delayed recall and Experiment 3: memory performance when instructed to remember with or without enactment before recall.

#### Experimental setup

The study did not require a specific experimental setup as all dolphins had to be tested in their home pools shared with other dolphins. As mentioned before, the dolphins were tested individually and kept apart by different trainers in separate locations of the pools, engaging with them throughout the test session, so that no mutual interference occurred. During work with a trainer, the dolphin stayed close to the trainer assigned to it for the session, at the edge of the pool. During waiting intervals it received little pieces of fish and gelatine from a bucket to make it stay nearby. Individual training or testing in interaction with various human trainers was a routine, daily procedure for the dolphins.

#### Experiment 1: Memory performance when anticipating immediate recall

##### Test criterion

After all animals had been trained to respond to the “repeat” command, they had to meet criterion before proceeding to the actual repeat test (see SI for details). They were tested in sessions consisting of 20 trials in which two commands altered pseudo-randomly. The first command was always an *“action command”* for one of the four trained actions, while the second command following ca. five seconds later was either a “*repeat command*” (12 “repeat” trials) or another “*action command”* for one of the four actions in pseudorandom and counterbalanced order (eight “action” trials). This procedure prevented the animals from learning to repeat the first command on every trial. Thus, after having received and executed the first “action command”, they had to retain or update their memory depending on whether they encountered a “repeat” or another “action” command. To reach criterion, the dolphin had to perform at least seven correct repeats out of 12 (58%) during a session.

##### Repeat baseline

To establish the baseline for the dolphins’ ability to repeat their own actions without delays, eight experimental sessions were completed after the dolphins had passed training criterion. For direct comparability, we followed the method by Torres Ortiz et al. 2022 (48) employed in parrots. Each session consisted of 26 trials comprising two commands per trial. Four of the trials were “single repeat” trials (15%), eight trials were “double repeat” trials (31%, i.e., 46% “repeats” in total) and 12 trials were “action trials” (46%). The remaining 2 trials (8 %) were just actions after the “double repeat” trials. In “single repeats” the previous action was repeated once. “Double repeats” were trials in which a “repeat” was followed by yet another “repeat command”, so that the subject had to carry out the same action three times consecutively (this served the purpose of testing the understanding of repeating the last action since repeating a “repeat” does not contain any information, see SM for the results). In “action trials”, an action was followed by the “action command” for another of the four actions, instead of a “repeat command”. The first response of the first “repeat” of a “double repeat” trial, was analysed as a single “repeat” trial for the results. All trials were double blind with the trainer wearing opaque sunglasses and the assistant communicating each command verbally to the trainer. The assistant also blew a whistle after each correct response so that the trainer knew when to reinforce the subject (and to signal to the subject whether the response was correct straight away).

##### Delayed repeat tests

To investigate for how long the dolphins could remember their own previous action when anticipating immediate recall (i.e., recall within five seconds), delays were introduced between the last action carried out upon command and the “*repeat command*” and also between “action” trials. The first trial started with a three-second delay and subsequently a staircase paradigm was used in which the delay increased three seconds after a correct “repeat” trial and remained the same during an “action” trial regardless of the outcome. If the animals performed the “repeat” trial incorrectly, the delay decreased by three seconds and was also held constant during an “action” trial, regardless of the outcome (each switch was defined as a reversal). With this method we aimed to determine the animals’ memory limit. We conducted 12 sessions with 16 trials per session. Each session had 12 “repeat” trials and 4 “action” trials intermixed randomly. The memory threshold was defined as the point at which an individual had reached at least six consecutive reversals during the 12 sessions and hence had not passed the next three-second delay.

Both the baseline and the repeat trials with delays were double blind with the trainer wearing opaque sunglasses and the assistant communicating each command verbally to the trainer following an R-script-generated randomized list (RStudio, version 1.1.383^75^). The assistant blew a whistle after each correct response so the trainers knew when to reward the dolphin. The delayed repeat test followed on straight after the repeat baseline without specific training for the delays between the “action” and the “repeat command”. During the delayed repeat test, the assistant used a computer and the R-script displayed the next command to be communicated to the trainer after the delay. The assistant entered if the response was correct or not, so that the delay for the following trial was automatically adjusted and visibly timed.

#### Experiment 2: Memory performance when anticipating delayed recall

##### General procedure and commands used

For both 15’-delay group and 3’-delay group dolphins, a trial started with the “*instruction to remember*” and two “*checks*” (see below), after which the actual delay commenced, and ended when the dolphin had displayed an action upon the final “*recall reques*t”. The “*instruction to remember”* consisted of a series of commands, starting with the “*remember command*” followed by an “*action command*” (signalling the respective action to be remembered) and ended with the “*go command*” upon which the dolphin performed the action it had been signalled as part of the instruction (see Fig 1A). The “*recall request*” consisted of the “*remember command*” and the “*go command*” *without* the “*action command*” in between (see Fig 2A). As part of a standardised procedure, two “*recall requests*” referred to as “*checks*” (see Fig 2A;) were implemented before the actual delay in order to ensure the dolphin had indeed paid attention and encoded the action to be remembered. The first “*check*” occurred after one minute and the second one after three minutes following the “*instruction to remember*”. If the dolphin displayed the correct action after both “*checks*”, the actual delay for the respective test phase (Table 1) ensued.

**Table 1.** Experimental test phases. Test phases of experiment 2 “Memory performance when anticipating delayed recall” and their descriptions for the 15’-delay group. Following the baseline, the 3’-delay group was tested directly in phase #3, and then received a single unexpected trial of phase #6. *The 16h delay phase involved an overnight break during which the dolphins slept. Sleep is known to support memory consolidation in other mammals, birds and invertebrate species^76,77^.

| Phase # | Delay | Distractions | Description |
| --- | --- | --- | --- |
| **Baseline** | ~ 10 seconds | No | “*instruction to remember*” at the beginning of a training session and “recall request” immediately after reinforcing with fish |
| **1** | ~ 15 minutes | No | “*instruction to remember*” at the beginning of a training session and “recall request” at the end |
| **2** | ~ 15 minutes | Yes ~ 41 | REMEMBER at the beginning of a training session and RECALL at the end |
| **3** | ~ 2 hours | No | REMEMBER at the end of a training session and RECALL at the beginning of the next training session |
| **4** | ~ 2 hours | Yes ~ 119 | REMEMBER at the beginning of a training session and RECALL at the beginning of the next training session |
| **5** | ~ 4 hours | Yes ~ 158 | REMEMBER at the beginning of a training session and RECALL at the end of the next training session |
| **6** | ~ 16 hours* | No but Sleep* | REMEMBER at the end of the last training session of the day and RECALL at the beginning of the first training session the following day (= overnight) |
| 7 | ~ 8 hours | Yes ~ 264 | REMEMBER at the beginning of the first training session of the day and RECALL at the beginning of the final training session at the end of the day |

At the end of the delay a trainer called the dolphin and instructed it to recall by the “*recall reques*t”. If the subject performed the correct action, which it had previously been asked to remember, the trainer blew a whistle, and right after, reinforced the dolphin with fish. If the subject displayed an incorrect action or did not respond at all, the trainer maintained a neutral position in front of the dolphin and waited for 3-5 seconds, during which it was waiting and attending to the trainer (“Least Reinforcement Scenario”, LRS). The neutral position indicated lack of reinforcement and was used to signal an incorrect response. The respective trial was counted as incorrect, and a new trial commenced with the next “*instruction to remember”* including another action command chosen from a randomised list of 6 possible actions (S1 Table). These 6 actions were chosen from the pool of trained behavioural actions each dolphin performed reliably on trainer command.

Throughout testing, all trainers wore sunglasses that prevented the dolphins from seeing the trainers’ eyes and kept their body still in order to prevent any unintended cueing of the animals. Additionally, around half of the test trials of the 15’-delay group were double-blind trials, in which the trainer who asked the dolphin to recall the remembered action after the delay was another person than the trainer who had instructed the dolphin to remember the action at the beginning of the trial. More double-blind trials were not possible due to limited trainer availability. The double-blind procedure ensured the trainer giving the “*recall request*” was naïve with respect to the action that the dolphin was supposed to remember and therefore could not inadvertently cue the dolphin to respond correctly^78^. The trainer who instructed the dolphin to remember stood at the back a few meters away and blew the whistle if the response was correct, so that the naive trainer knew when to reinforce the dolphin.

##### Experimental groups

Experiment 1 was run with two experimental groups, a 15’-delay group that anticipated delayed recall in the more distant future and a 3’-delay group that had never experienced recall later than three minutes, thus after a not very long delay. During their training, the 15’-delay group had experienced increasingly longer delays before the “recall request” amounting up to 15 minutes and had reached training criterion with a 15 minute delay. The 3’-delay group in contrast, experienced delays only up to three minutes during their training and reached training criterion for this three minute delay (see SI for details).

##### Baseline and test phases

After having reached the training criterion (remembering correctly in 10 out of 18 trials; see SI for further detail), all dolphins were first tested in a baseline and then took part in several test phases (Table 1) consisting of 18 trials each. The 15’-delay group underwent 7 test phases (Table 1) with gradually increasing delay duration (15 minutes with and without distraction, 2 hours with and without distraction, 4 hours, 8 hours and 16 hours), whereas the 3’-delay group was tested straight away in the 2h test phase after the baseline and finally received one single unexpected 16/17 hours delay recall test. The baseline consisted of 18 trials in which no actual delay occurred, i.e., the “*recall request*” was given straight after the “*instruction to remember*”. This still involved a brief lag of ∼ 10 seconds due to the time needed to reinforce the dolphin after performing the action to be remembered following the first “*go command*”. The baseline was collected to control for mistakes occurring naturally due to a lack of attention or other parameters. Additional test phases with delays longer than 16h could not be tested due to time constraints of the study.

The 15’-delay group was instructed to remember one out of six actions and the 3’-delay group with one out of four actions (selected from each dolphin’s respective repertoire of trained behaviours it performed on command, see SI and S1 Table) in pseudorandom and counterbalanced order. For both groups a minimum of ten trials (At least 55% of the 18 trials, see explanation above) was double-blind.

##### Examining the effect of distractions

In order to judge how distraction (see definition below) during a delay interval between the “*instruction to remember*” and “*request to recall*” would affect the dolphins’ performance, we tested the individuals in the 15’-delay group with and without distractions for the 15-minute delay and for the 2h delay. The remaining test phases were either with or without distractions (see Table 1). In the trials without distraction, the dolphin was instructed to remember an action and then could swim freely but without interacting with any human trainer throughout the predetermined delay period. At the end of the delay, the dolphin was called by the trainer and requested to recall the remembered action. In the trials with distraction in contrast, the dolphin engaged in different activities with human trainers throughout the delay period (and also moved freely in the pool with the other dolphins). Every action the dolphin was asked to perform upon hand command following the “*instruction to remember”*, was considered as a “distraction”. For example, if a dolphin performed 12 different actions associated with hand signals, the number of distractions during that delay was considered as 12. During phases with distractions, the trainer either simply continued with a characteristic training session, further training session(s) with the same or different trainers took place later on, a medical procedure was carried out (i.e., blood sample, ultrasound session) or the dolphin participated in one of the dolphin programs of the facility. All those sessions were composed by different actions (jumps, position for a blood sample, among many others). Those actions that the dolphin was asked to carried out were defined as distractions (one action = one distraction). The number of actions the dolphin was asked to carry out during the interval of a test trial was counted by one of the trainers throughout four training sessions occurring during the delay of each phase. This allowed us to calculate the number of actions per minute to estimate the approximate number of actions for the remaining trials in each phase with distractions.

#### Experiment 3: memory performance comparing previously enacted versus gesturally encoded actions

The 3’-delay group dolphins underwent a further memory test. They were requested to remember the trainer’s hand gesture for an action the dolphin was supposed to carry out (as soon as the “*go command*” was given), but without letting the dolphins self-perform the behaviour. Hence, for this experiment, the “*remember command”* was implemented with a delay introduced before the “*go command*”, i.e., in the following manner: (“*remember command” + “action command”* + delay + “*go command”*). The two dolphins were tested with a 10-second delay and a 60-second delay. Each delay condition comprised 8 trials throughout which they had to remember the four actions in a pseudorandomized and counterbalanced manner. The data was directly compared to the previously collected baseline ∼10 second (± 2 seconds) and the one-minute delay from Experiment 2.

### Quantification and statistical analysis

All analyses were performed using the *rethinking* ^79^ package in R ^75^. For the average performance we report the average performance and 89% posterior interval (PI) on the probability scale (percentage correct). For the effects of the continuous, binomial and ordinal variables we report the average and 89% PI for the slopes on the log-odds scale. In the figures we plot the average predictions as well as 20 samples from the posterior. When comparing between two conditions we report the average and 89% PI for the contrast of the intercepts for these two conditions on the log-odds scale. A contrast consists of the difference between the two posterior distributions of the two conditions. If this contrast does not or only very little overlaps with zero, it means there is a clear difference between conditions.

All models were fitted using Stan ^80^ with the ulam ^79^ and cmdstanr ^81^ interface, with 4 chains, 4000-8000 iterations, 500 of which were the warm-up phase. We monitored all diagnostics and ensured divergent transitions were never above 0.1%. We also monitored rhat values and ensured they were never more than 0.01 above or below 1.

#### Performance in experiment 1 when anticipating immediate recall

##### Performance in the repeat baseline

The following model was used to investigate individual performance in the repeat baseline:

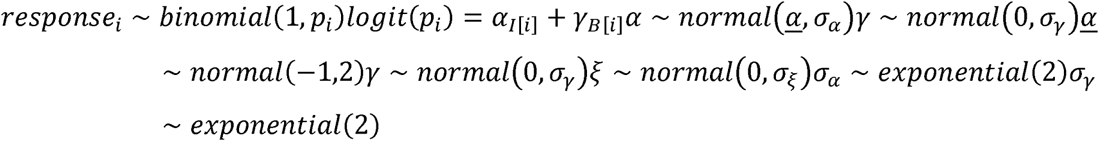

Where I = individual and B = behaviour.

##### Performance in the delayed “repeat” test

To model the effect of delay on the performance during the delayed “repeat” test, we used a similar model, where log(time) was included as an additional variable:

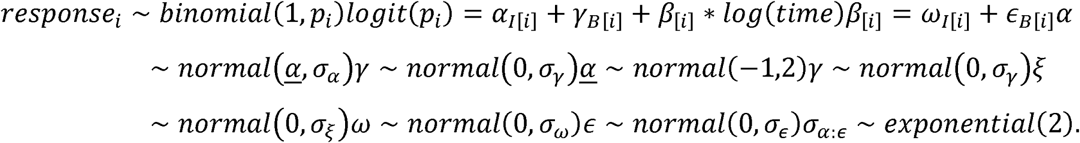

Where I = individual and B = behaviour.

#### Performance in experiment 2 when anticipating delayed recall

Since the experiment was conducted in discrete phases, with each phase representing a discrete increase in difficulty, we treated the phase as an ordered categorical variable. We used a similar Bayesian logistic model, including animal, behaviour, double blind, date, trainer marking the behaviour to be remembered and trainer requesting the dolphin to recall as varying effects on the intercept. We also included the normalized natural logarithm of the number distractions and the phase as predictors. We included varying effects for animal on the slopes, to allow individuals to respond differently to these challenges.

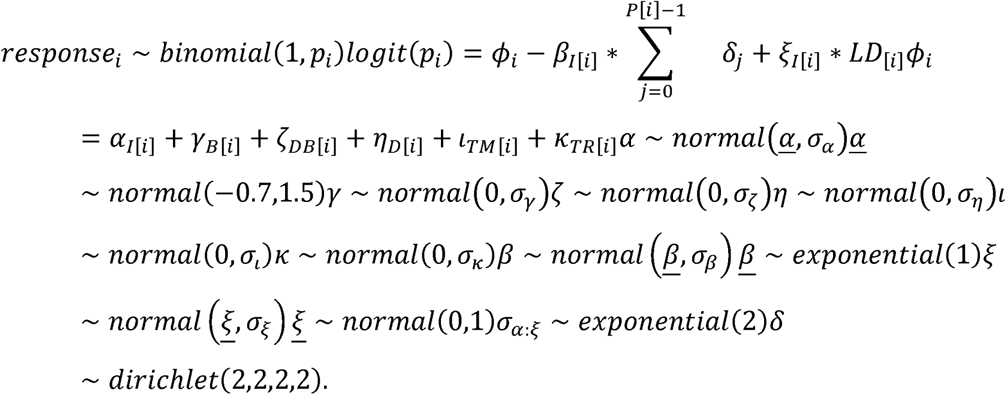

Where I = individual, B = behaviour, DB = double blind, D = date, TM = trainer mark, TR = trainer recall and LD = normalised log number of distractions. We restricted the slope of the phase to negative values, but allowed positive values for the slope of the distractions.

To compare the performance between the 3’-delay dolphins (without explicit training for delays) and 15’-delay dolphins (with explicit training for delays), we used a similar model, including the experimental group as varying effect:

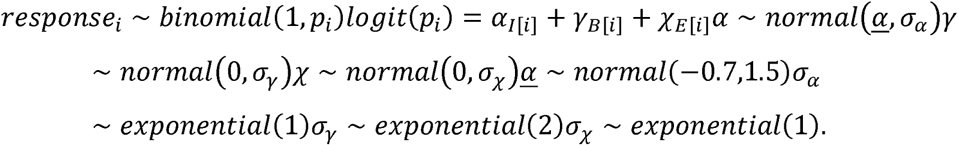

Where I = individual, B = behaviour and E = experimental group. The models for the 3 minute and 2 hour delay experiments were the same.

#### Performance in experiment 3 comparing the memory of previously enacted versus gesturally encoded actions

To model the effect of remembering an action not self-performed during the encoding (enactment effect), we ran a model that included experimental group (enacting vs not enacting) as varying effect and also included the phase (10 seconds vs 60 seconds) interacting with the experimental group as varying effect (meaning that we included an off-set for each combination of experimental group and phase - four off-sets in total). We also included a varying effect for individual. The model had the following structure:

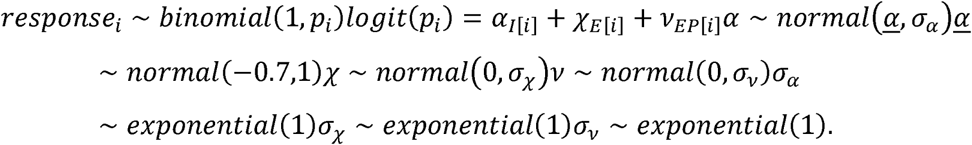

Where I = individual, E = experimental group and EP = experimental group and phase.

#### Ethics statement

All applicable international, national and/or institutional guidelines for the care and use of animals were followed. The animals from Dolphin Adventure were kept and trained under permits from Mexican Secretariat of Environment and Natural Resources [permit: INE/CITES/DGVS-EF-P-0033-NAY/00 (PIMVS)]. In accordance with the German Animal Welfare Act of 25th May 1998, Section V, Article 7 and the Spanish Animal Welfare Act 32/2007 of 7th November 2007, Preliminary Title, Article 3, the study was classified as non-animal experiment and did not require any approval from a relevant body.

All subjects from both facilities participated in the study on a voluntary basis. The dolphins were not food-deprived and could choose to come to the experimenter or leave the session anytime. Their full daily diet was given regardless of participation.

## Acknowledgments

We thank Dolphin Adventures for their support, access to the dolphins and the trainers” time, particularly Lucio Conti, Benjamin Doshner, Juan Carlos Chavez and Wayne Phillips. We thank Loro Parque and its president, Mr. Wolfgang Kiessling for their support, the access to the dolphins and trainer support. We thank the Loro Parque Fundación and its president Mr. Christoph Kiessling for their collaboration and the staff of the Loro Parque Fundación, the animal caretakers and the veterinary department for their constant support. Lastly, we are grateful to Christian Leyva Cortes and Javier Luis Lopez for assisting in part of the data collection and to Manfred Gahr and Jason Bruck for comments to improve the manuscript.

## Author contributions

S.T.O. and A.v.B. designed and supervised the entire project. S.T.O., A.M.G., A.H.S., and C.U.R. trained the animals and collected the data. S.T.O., and S.Q.S. analysed the data and prepared the figures. S.T.O., S.Q.S., M.O. and A.v.B. wrote the first draft, and S.T.O and A.v.B. wrote the final manuscript supported by S.Q.S. and M.O. All authors were involved in the data interpretation and commented on and approved of the manuscript.

