## Supplemental Information for "Awareness of the future: Dolphins know when they need to remember for the future"

**Supporting information**

**Previous behavioural training and training criterion**

Over many years, the five dolphins in the 15’-delay group (Dolphin adventure) and the three dolphins in the 3’-delay group (Loro Parque) had been behaviourally trained to perform specific actions on command (human hand signals) using operant conditioning with positive reinforcement ^81,82^. When the dolphins performed the correct action on command, the trainer blew a whistle instantly signalling to the dolphin that the behaviour was correct and then reinforced it with fish. If the dolphin’s response was incorrect, the trainer did not blow the whistle but implemented a 3-5 second break during which the dolphin was kept waiting to indicate that the displayed behaviour was incorrect.

For the 15’-delay group dolphins, six behaviours from each dolphin’s repertoire of previously trained actions were chosen for the experiment, which the dolphins performed reliably on specific commands. For the subsequently tested Loro Parque dolphins, which had been first trained for Experiment 1, then served as 3’-delay group for Experiment 2 and finally were used for Experiment 3, four behaviours from their repertoires were chosen. The selected trained actions involved comparable effort and were firmly established (i.e., the chosen behaviours had been part of the dolphin’s training sessions during several years; S1 Table). After phase 1, “splash” was replaced with “go down” for the subject Karina, due to mouth injuries.

**S1 Table. Experimental actions**. Description of the trained actions selected from the dolphins’ repertoire for this study and list of which dolphins performed which actions.

| Action | Description | Subject |
| --- | --- | --- |
| Clap | Dolphin in vertical position with head and pectoral flippers out of the water moves its pectoral flippers back and forward rapidly | Exp. 2 15’-delay group: Karina, Nemo, Nouba, Lluvia, Eva  Exp. 2 3’-delay group: Achille, Ulisse  Exp. 1 & 3: Achille, Ulisse, Clara |
| Spin | Dolphin in vertical position with head out of the water turns around itself | Exp. 2 15’-delay group: Karina, Nemo, Nouba, Lluvia, Eva  Exp. 2 3’-delay group: Achille, Ulisse Exp. 1 & 3: Achille, Ulisse, Clara |
| Tail wave | Dolphin in vertical position but upside down with tail out of the water moves the tail back and forward repeatedly | Exp. 2 15’-delay group: Karina, Nemo, Nouba, Lluvia, Eva  Exp. 2 3’-delay group: Achille, Ulisse Exp. 1 & 3: Achille, Ulisse, Clara |
| Jump | Dolphin jumps out of the water with its entire body | Exp. 2 15’-delay group: Karina, Nemo, Nouba, Lluvia, Eva |
| Splash | Dolphin in vertical position with head out of the water squirts/spits water out of its mouth | Exp. 2 15’-delay group: Karina (only phase 1), Nemo, Nouba, Lluvia, Eva |
| Sing | Dolphin in vertical position with head out of the water produces vocalisations | Exp. 2 15’-delay group: Karina, Nemo, Nouba, Lluvia, Eva |
| Go down | Dolphin sinks to the bottom of the pool until it lies there horizontally on its belly | Exp. 2 15’-delay group: Karina |
| Belly up | Dolphin turns its body upside down with its belly showing on the water surface | Exp. 2 3’-delay group: Achille, Ulisse  Exp. 1 & 3: Achille, Ulisse, Clara |

**Specific training for Experiment 1 (Memory performance when anticipating immediate recall)**

***“Repeat command” training (Loro Parque dolphins, 3’-delay group)***

The three dolphins (S2, S3 Table) were trained on the “*repeat command*” using the identical training steps that were implemented in parrots by Torres Ortiz et al. ^23^:

1. The trainer gave the respective command (hand gesture) for a specific action (e.g., spin) five times consecutively letting the subject perform it in succession and right after introducing the hand gesture for the “*repeat command*”. Usually, the subject would do the previous action (e.g., spin) because of being in the “flow”. If this was not the case, the trainer “helped” by pairing the “*action command*” with the “*repeat command*”.
2. Once the dolphin performed the first instructed action upon “*repeat command*” without “help” cues, the trainer instructed the subject to perform another action five times consecutively by giving the respective action command (i.e., hand gesture for a different specific trained action, e.g., belly up) five times, and right after, gave the “*repeat command”*. Training sessions continued until the dolphins repeated those two actions reliably discriminating between the two actions well.
3. Next, the trainer introduced a third action and asked the dolphin to repeat it, to test if the dolphin was able to generalise the *“repeat command*” to novel actions based on the previous training. If the dolphin did not respond correctly, training followed the previous training steps (1 and 2). Training sessions continued until the dolphin had reliably repeated the three randomly intermixed actions.
4. Finally, the trainer introduced the last action and tested once more if the dolphins applied the “repeat” concept to another action. If the animal failed, the trainer again followed the previous training steps (1 and 2) until the animal successfully repeated the fourth action on “*repeat command*”. Training sessions were performed until the animal reliably repeated in training sessions in which the four trained actions were intermixed pseudo-randomly.
5. The training continued until the subject passed criterion, which was to repeat seven times correctly out of 12 repeat trials intermixing the 4 actions and 16 additionally intermixed action trials (requesting the subject to perform an action on command without following “repeat” (with a 0.25% chance level).

**S2 Table. Experimental gestural commands**. Description of the hand signals trained and used in our study in addition to the previously trained action commands.

| Command | Description | Hand gesture used |
| --- | --- | --- |
| “*Remember command*” | Command used to instruct the dolphin to remember the following content. The “*remember command*” is followed by the “*action command*” and the “*go command*” and all three together comprise the “***instruction to remember”*** (Fig 2A).  Combined with just the “*go command*” on the other hand, it constitutes the  **“*recall request*”** (Fig 2A). | Both index fingers pointing at the dolphin |
| *“Go*  *command”* | Command used to let the dolphin know that he can proceed to perform the action that the trainer requested the dolphin to remember or recall. | Both fists hitting each other vertically |
| *“Repeat”* | Command used to tell the dolphin to repeat the previously performed behaviour | Both hands spinning around each other in circles |

**S3 Table. Experimental timeline.** Information on the time period in which the study was conducted with each subject.

| **Individual** | **Group** | **Experiment** | **Type** | **Start date** | **End date** |
| --- | --- | --- | --- | --- | --- |
| Nemo | 15’-delay condition | Exp. 2: Prospective encoding | Training | 22/04/2020 | 21/06/2020 |
|  |  | Exp. 2: Prospective encoding | Testing | 22/06/2020 | 21/10/2020 |
| Karina | 15’-delay condition | Exp. 2: Prospective encoding | Training | 22/04/2020 | 21/06/2020 |
|  |  | Exp. 2: Prospective encoding | Testing | 22/06/2020 | 02/11/2020 |
| Eva | 15’-delay condition | Exp. 2: Prospective encoding | Training | 22/04/2020 | 21/06/2020 |
|  |  | Exp. 2: Prospective encoding | Testing | 22/06/2020 | 22/10/2020 |
| Nouba | 15’-delay condition | Exp. 2: Prospective encoding | Training | 22/04/2020 | 21/06/2020 |
|  |  | Exp. 2: Prospective encoding | Testing | 22/06/2020 | 15/10/2020 |
| Lluvia | 15’-delay condition | Exp. 2: Prospective encoding | Training | 22/04/2020 | 21/06/2020 |
|  |  | Exp. 2: Prospective encoding | Testing | 22/06/2020 | 28/10/2020 |
| Clara | 0’-delay condition | Exp. 1: Repeat | Training | 17/03/2021 | 30/09/2021 |
|  |  | Exp. 1: Repeat | Testing | 01/10/2021 | 08/10/2021 |
| Achille | 0’- and 3’-delay condition | Exp. 1: Repeat | Training | 07/10/2021 | 15/01/2022 |
|  |  | Exp. 1: Repeat | Testing | 18/01/2022 | 28/01/2022 |
|  |  | Exp. 2: Prospective encoding | Training | 16/05/2022 | 14/06/2022 |
|  |  | Exp. 2: Prospective encoding | Testing | 15/06/2022 | 27/08/2022 |
|  |  | Exp. 3: Reenactment test | Testing | 29/09/2022 | 14/10/2022 |
| Ulisse | 0’- and 3’-delay condition | Exp. 1: Repeat | Training | 13/09/2021 | 06/12/2021 |
|  |  | Exp. 1: Repeat | Testing | 10/12/2021 | 08/01/2022 |
|  |  | Exp. 2: Prospective encoding | Training | 16/05/2022 | 14/06/2022 |
|  |  | Exp. 2: Prospective encoding | Testing | 15/06/2022 | 18/09/2022 |
|  |  | Exp. 3: Reenactment test | Testing | 29/09/2022 | 14/10/2022 |

**Specific training for Experiment 2 (Memory performance when anticipating delayed recall)**

The commands that instructed the dolphins to remember, recall or repeat a particular action were established as described below (S2 Table).

**“Remember command” training (Puerto Vallarta dolphins, 15’-delay group)**

*Description of the command series to be trained:*

Throughout the training to acquire the “remember command”, the dolphins learned a series of commands consisting of three different hand signals (“*remember command”* = remember actively + “*action command*” *=* behaviour to be performed and remembered + “*go command”* = perform now), i.e., the “*instruction to remember*”. Following the “*go command”,* the subject had to perform the action indicated by the respective “*action command*” and was rewarded if it enacted the indicated action correctly.

After this instruction, we asked the dolphins to recall and perform the remembered behaviour (“*recall request*”), hence the action indicated within the previous *“instruction to remember*” The series of commands also consisted of “*remember command”* and “*go command”* but omitted the “*action command*”. Now, the “*go command”* meant that the dolphin had to perform the action that it had been instructed to remember even though the respective “*action command*” was not presented.

As described in the methods, the “recall request” was repeated twice more, following an interval of one minute and 3 minutes respectively (= one- and three-minute “checks”) in order to ensure the dolphin had paid attention. If the dolphin passed both “checks”, the respective delay ensued, followed by the actual test “*recall request*”. If a subject failed, the “checks” the procedure was restarted with a new “*instruction to remember*”.

*Gradual training process*

In order to train the “*instruction to remember*”, the dolphin was first introduced to the hand signal of the “*remember command*” (S2 Table) and reinforced for paying attention to the trainer. After this, the subject was instructed to carry out an action from their repertoire of trained actions, which it had known for years, by giving the “*action command*” specifying that particular action following the “*remember command*”. Yet, the dolphin was prevented from performing that action straight away (by making them touch the palm of the trainer’s hand) in order to teach it to wait for a while before performing the action when the “*action command*” was given following the “*remember command*”. Once the dolphin had learned not to perform the action straight after the “*remember command*” + “*action command*”, the “*go command*” was introduced and, as the next step, the dolphin was trained to perform the action only after the “*go command*” had been given. Once the dolphin reliably waited until the “*go command*” was given to perform the specific requested action, i.e., when it had learned the full series of commands (“*remember command” + “action command” + “go command”*), the trainer started introducing the “*recall request*”, i.e., the “*remember command*” and “*go command*” without giving the previous “*action command*” again, straight after the “*instruction to remembe*r” and thus requested the dolphin to perform the previous action again. This entire procedure was repeated with all the 6 actions chosen for the study (S1 Table) introduced one after the other.

After successful learning of the “recall request” and establishing the “*recall request*” after one minute and three minutes (“*checks*”) after the “*instruction to remember*”, the delay in between the two was gradually introduced. First, the time between the “three-minute check” (i.e., the last “*recall request*” as part of the remembering process) and the actual final “*recall request*” was increased in small steps until a 15-30 second delay was reached for each participating dolphin. Next, distracting activities were introduced, i.e., the dolphins were instructed to perform other behaviours during the delay phase before receiving the final “*recall request*”. The trainers started with requesting simple behaviours like swimming around. Finally, the trainer instructed the dolphin to carry out various actions, including actions that were similar to the one they had been requested to remember during the delay.

Each 15’-delay group dolphin was trained with six actions chosen from their trained behavioural repertoire (S1 Table) until a delay of 15 min was reached. The training of the “*instruction to remember*” and “*recall request*” for the experiment started on the 22^nd^ of April of 2020. Testing started when dolphins had reached a criterion of 10 correct trials from a total of 18 trials (55% correct) with a delay of 15 minutes without distractions occurring during the delay. The first testing session started on the 22^nd^ of June 2020. One to two trials were carried out per day, with at least a two-hour inter-trial period. The experiment ended the 2^nd^ of March of 2021.

**Specific training for Experiment 2 and 3 (Memory performance when anticipating delayed recall and enactment effect)**

***“Remember command” training for the Loro Parque dolphins (3’-delay group)***

Two dolphins (Achille and Ulisse) constituted the 3’-delay group for Experiment 2. The training of the 3’-delay group started the 16^th^ of May 2022 (S3 Table). The training steps were the same as for the 15’-delay group, except that the 3’-delay group dolphins had not been trained with delays incrementally increasing in duration until 15 minutes between “instruction to remember” an action and “recall request” were reached. Instead, they were trained for 20 days continuously with delays not exceeding three minutes. Then from the 9^th^ of June 2022, the trainers started to introduce distracting actions between the “*instruction to remember*” the action and the “*recall request*” of the action that the dolphin had to remember. However, the delays between memorization and recall still never exceeded three minutes. Data collection with the 3’-delay dolphins started the 15^th^ of June of 2022 (S3 Table). Testing started when dolphins had reached a criterion of 10 correct trials from a total of 18 trials (55% correct) with a delay of max. three minutes without distractions occurring during the delay. The first testing session started on the 16^th^ of May of 2022. One to two trials were carried out per day, with at least a two-hour inter-trial period. The experiment ended the 18^th^ of September of 2022.

**Supplementary figures and videos**

**S1 Fig. Dolphin adventure first facility.** Aerial view of the dolphin enclosure at Dolphin Adventure, Puerto Vallarta where the four females, Karina, Eva, Nouba and Lluvia were housed together with 12 other dolphins. This facility is inland with concrete-built pools.

**S2 Fig. Dolphin adventure second facility.** Aerial view of the dolphin enclosure at Dolphin Adventure, Puerto Vallarta where the male, Nemo, was housed together with six other dolphins. This facility is in the harbour and the pool walls are made of sailcloth.

**S3 Fig. Loro Parque facility.** Aerial view of the dolphin enclosure at Loro Parque: Animal Embassy, Tenerife where the 3’-delay group dolphins, Achille, Ulisse and Clara were housed. This facility is inland with concrete-built pools.

**S1 Video. Video of one experimental session for experiment 1: memory performance when anticipating immediate recall of own actions after short delays**. The experimenter in front of the dolphins is wearing blinded sunglasses, the experimenter holding the computer is communicating which action command should be given to the dolphin and the trainer on the side is blowing the whistle to let the dolphin and the blinded experimenter if the response is correct.

**S2 Video. Video of one of the experimental sessions for experiment 2: memory performance when anticipating delayed recall.** The “*instruction to remember*” is given by one experimenter to encode the action to be remembered. After the delay, a second experimenter naïve of what the dolphin was told to remember provides the “*recall request*” following which the dolphin should reenact the specific action that it was told to remember.

**S3 Video. Video of the two types of sessions for experiment 3: memory performance comparing the memory of previously enacted versus gesturally encoded actions.** The experimenter is giving the dolphin the “*instruction to remember*” without the “*go command*” so that the dolphin is told what to remember but is not enacting the memory content until a 10 second delay has passed. The second half of the video illustrates the same experiment but with a 60 second delay from the “*instruction to remember*” until the “*go command*” is given.
