## Supplementary figures and images for "Awareness of the future: Dolphins know when they need to remember for the future"

### Supplemental Figure 1

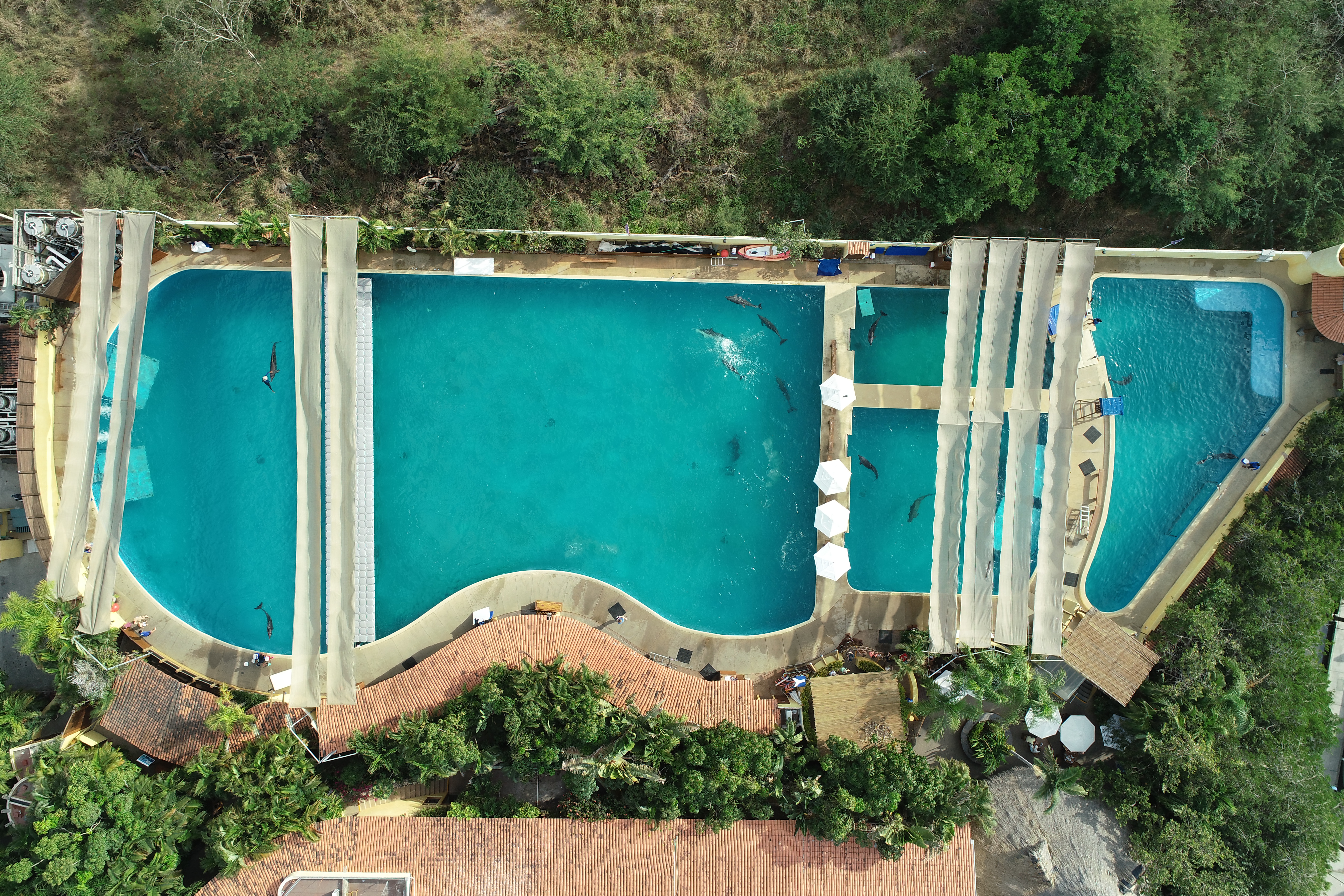

### Supplemental Figure 2

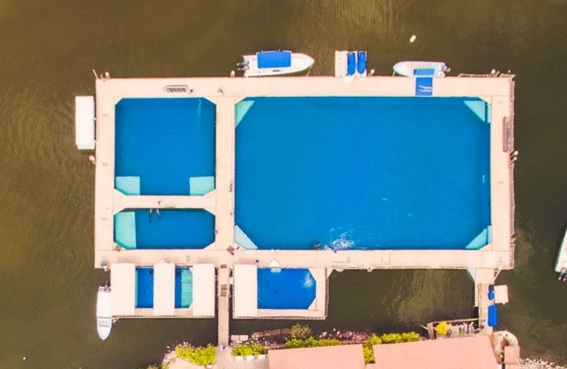

### Supplemental Figure 3

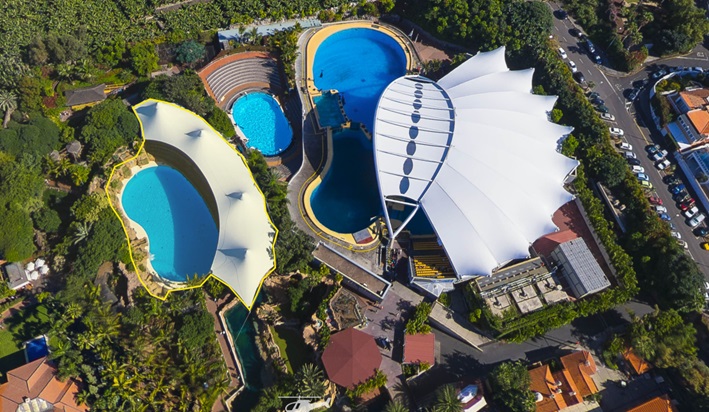
